# Deciphering Reductive Dehalogenase Specificity Through Targeted Mutagenesis of Chloroalkane Reductases

**DOI:** 10.1101/2024.07.29.605694

**Authors:** Katherine Picott, Connor Bowers, Elizabeth A. Edwards

## Abstract

Reductive dehalogenases (RDases) are essential in the anaerobic degradation of various organohalide contaminants. This family of enzymes has broad sequence diversity, but high structural conservation. There have been few studies assessing how RDase peptide sequences affect their substrate selectivity. Here we focus on two chloroalkane RDases, CfrA and DcrA, which have 95% protein sequence identity but have diverged to hold distinct substrate preferences. CfrA will dechlorinate chloroform and 1,1,1-trichloroethane, whilst DcrA will dechlorinate 1,1-dichloroethane. We mutated several residues in the active site of CfrA to investigate a change in substrate preference and to identify which wild-type residues contribute the most to substrate specialization. We determined that no individual residue solely dictates substrate discrimination, but both Y80W and F125W mutations were needed to force CfrA to prefer 1,1-dichloroethane as a substrate. This double mutation also altered the transformation pathway of 1,1,2-trichloroethane from hydrogenolysis (forms 1,2-dichloroethane) to dihaloelimination (forms vinyl chloride). We use predictive protein models and substrate docking to predict what interactions are made between the enzyme and substrate to aid in selection. The residues of significance identified in this study are consistent with those identified from chloroethene RDases, suggesting residue locations with a particularly high impact on activity.

**Importance:** Reductive dehalogenases play an integral role in the removal of chlorinated solvents from the environment. These enzymes have specificity towards different chlorinated compounds, and it is known that small natural changes in their peptide sequence can change their activity drastically. How these specific sequence variations influence activity is largely unknown. In this study, we demonstrate that mutating a few residues within the active site of CfrA—a chloroform and trichloroethane-specific dehalogenase—changes its substrate preference to dichloroethane. We determine that only two mutations are needed to disrupt the native activity, underscoring the nuances in substrate-structure relationships in reductive dehalogenases. Though we are still far from predicting function from the sequence, this knowledge can give some insight into engineering reductive dehalogenases for new target contaminants.

## Introduction

Organohalide-respiring bacteria (OHRB) are well-known for their role in the bioremediation of chlorinated solvents such as chloroethenes and chloroethanes, many of which are on the priority substance list for toxic compounds (1). OHRB utilize organohalides as their terminal electron acceptor, which are reduced by membrane-bound reductive dehalogenases (RDases) in the electron transport chain (2). RDases catalyze the cleavage of halogen-carbon bonds during the reduction reaction (2). This cleavage can occur via hydrogenolysis, where the halogen is replaced with a proton, or through dihaloelimination (a β-elimination reaction) (2). RDases require a cobamide and two [4Fe-4S] clusters as cofactors for catalysis (3, 4), all of which participate in the reduction of the substrate.

RDases are not unique to OHRB, but are a vast family of enzymes that are found in many different bacteria and even archaea (5, 6). Accordingly, there is high variation in protein sequences; for example, characterized perchloroethene (PCE) dechlorinating RDases from different organisms can have less than 20% pairwise amino acid sequence identity (PID) with each other but maintain the same function (7–13). This high variation makes predicting RDase activity from sequence alone very difficult. An organizational system, the ortholog group (OG) system, was developed to help keep track of functionally characterized RDases and relate new sequences to potential substrates based on sequence similarity (14, 15). RDases are grouped into OGs based on having >90% PID between all members of the group (14, 15). While these OGs do indeed group RDases that have functions on the same class of compound (i.e. chloroethene versus chloroethane), the exact substrate specificity of RDases is even more discriminating.

OG 97 contains seven characterized chloroalkane reductases (all have >93% PID) and have shown varying substrate preferences (16–21). The starkest difference in selectivity comes from CfrA and DcrA, two enzymes that were identified from the same mixed culture enriched on 1,1,1-trichloroethane (TCA) (16, 22). CfrA and DcrA are expressed by the related organisms *Dehalobacter* sp. CF and *Dehalobacter* sp. DCA (23), respectively. Rather than competing for the same substrates, CfrA became specialized in 1,1,1-TCA and chloroform (CF) dechlorination, while DcrA primarily dechlorinates 1,1-dichloroethane (DCA) (16). Further, when given 1,1,2-TCA as a substrate CfrA and DcrA will undergo two different transformation mechanisms (24). CfrA predominantly performs hydrogenolysis to 1,2-DCA and DcrA prefers dihaloelimination to vinyl chloride (VC) (24). This natural evolution of these enzymes with high sequence identity yet contrasting function is an excellent opportunity to identify key residues that influence substrate preferences and gives a glimpse into the target areas for positive selection.

There have only been three RDase crystal structures solved to date: PceA from *Sulfurospirillum multivorans* (referred to as PceA_S_ from here on) (3), the cytosolic NpRdhA from *Nitrareductor pacificus* pht-3B (4), and most recently PceA from *Desulfitobacterium hafniense* TCE1 (PDB IDs: 4UQU, 4RAS, and 8Q4H, respectively) (25). The central fold of RDases, made up of α-helices and β-sheets, is highly conserved and researchers have been able to glean information on potentially important and influential residues; the residues we discuss here are numbered with respect to PceA_S_. Two active site amino acids— Tyr246 and Arg305—were identified in the PceA_S_ crystal structure and were subsequently demonstrated to be essential catalytic residues through mutagenesis (3, 4, 26). These two residues are highly conserved amongst RDases and are predicted to be involved in the final protonation step of dehalogenation aided by residue Asn272 (3, 4, 26). Additional residues have either been proposed to be involved in substrate selection based on *in silico* analysis or through mutagenesis studies (26–28). One mutagenesis study found several residues—corresponding to Trp96, Thr242, Trp376 in PceA_S_—in the active sites of both PceA from *D. hafniense* Y51 (referred to as PceA_D_ going forward) and DcaA from *D. dichloroeliminans* DCA1 (88% PID between PceA_D_ and DcaA) that modify the size of the substrate preferred by the enzymes when mutated (28). Additionally, Phe38 lines the active site and is proposed to help orient the substrate for catalysis, though this has never been tested (26, 27). These residues are all intriguing targets to assess substrate preference divergence between CfrA and DcrA. Table 1 compares these corresponding residues, analyzed through a multiple sequence alignment using MAFFT v7.017 and structural analysis using PyMOL (29, 30), along with descriptions of potential functions based on literature.

**Table 1.** Residues of interest identified in previous studies on PceA and DcaA and their putative roles.

| PceAs<br>(*S. multivorans*) | DcaA<br>(*D. dichloroeliminans*<br>DCA1)<sup>a</sup> | CfrA<br>(*Dehalobacter* sp.<br>CF) | DcrA<br>(*Dehalobacter*<br>sp. DCA) | Proposed Role/DcaA Mutagenesis<br>Observations | Ref |
| --- | --- | --- | --- | --- | --- |
| F38 | F31 | Y80 | W80 | Lines substrate channel and active site to orient substrate, no mutagenesis performed | (26, 27) |
| W96 | W118 | F125 | W125 | Active site formation, W118F lost activity | (26, 28) |
| T242 | T294<sup>b</sup> | Y256 | Y256 | Substrate interaction, T294V shifted activity towards smaller substrates | (25, 27, 28) |
| Y246 | Y298 | C260 | Y260 | Catalytic residue and suggested proton donor or stabilizing residue, Y298F eliminated hydrogenolysis activity but enhanced dihaloelimination | (3, 4, 26–28, 31, 32) |
| N272 | N324 | S286 | A286 | Stabilizes catalytic residues, no mutagenesis performed | (3, 26–28, 31, 32) |
| R305 | R357 | R319 | R319 | Catalytic residue suggested as a proton donor or stabilizing residue, R357X decreased activity on chloroethenes but increased on tribromoethene and dihaloelimination | (3, 26–28, 31, 32) |
| W376 | W432 | M391 | W391 | Lines substrate channel and active site to orient substrate, W432F decreased activity on chloroethenes but increased on tribromoethene and dihaloelimination | (26, 28) |

<sup>a</sup> Residues are consistent in PceA<sub>D</sub> from *D. hafniense* Y51, <sup>b</sup> V294 in PceA<sub>D</sub>

In this study, our aim is to broaden our knowledge of RDase-substrate interactions beyond haloethenes. By investigating natural active site variation between the chloroalkane-reductases CfrA and DcrA, we seek to expand our understanding of how RDases fine-tune their specificity. Through site-directed mutagenesis of select residues, we endeavour to identify the amino acid positions that are most influential in determining substrate selectivity and preferred transformation pathways.

## Materials and Methods

### Chemicals and Other Materials

Unless otherwise stated, all chemicals, BugBuster^®^, and filter tubes were purchased from Millipore Sigma (Burlington, MA, USA). All growth media, antibiotics, and induction agents were purchased from BioShop Canada Inc. (Burlington, ON, Canada). PCR mix, restriction enzymes, and Gibson assembly mix were purchased from New England Biolabs Ltd. (Ipswich MA, USA). Bradford reagent was purchased from Bio-Rad Laboratories Ltd. (Hercules, CA, USA). All gasses were supplied from Linde Canada Inc. (Mississauga, ON, Canada). A Coy vinyl anaerobic chamber (Coy Laboratory Products Inc., Grass Lake, MI, USA) and a gas supply of 10% CO_2_/10% H_2_/Bal. N_2_ (v/v) was used to maintain an anaerobic atmosphere, this is referred to as a glovebox throughout. Gas chromatography materials were purchased from Agilent Technologies (Santa Clara, CA, USA). All genes were confirmed by Sanger sequencing at The Centre for Applied Genomics (TCAG; SickKids, Toronto, ON).

### Sequence and Structure Comparison

The amino acid sequences of all proteins in OG 97 (CfrA, DcrA, TmrA, AcdA, CtrA, ThmA, and RdhA D8M_v2_40029) were aligned using MAFFT v7.017 plugin in Geneious v8.1.9 with default settings (29). Predictive protein models were made for CfrA, DcrA, TmrA, and AcdA using AlphaFold2 (33). Each non-conserved residue in the alignment was visualized in the protein models using PyMOL v2.3.4 to determine their proximity to the putative substrate binding site and their likelihood to contribute to activity differences observed between the enzymes (30). Five variable residues at positions 80, 125, 260, 388, and 391 were identified in the active site and were pursued for mutagenesis studies. Residues 80, 260, 388, and 391 were also noted as potentially important residues in the isotope fractionation study comparing TmrA, CfrA, and AcdA (34).

### Plasmid Construction

Plasmids for CfrA and DcrA were constructed previously (24). Genes for the mutants CfrA-5M, CfrA-3M, and DcrA-5M were codon optimized and purchased from Twist Bioscience (San Francisco, CA, USA) and cloned into *p15TV-L* plasmid as described previously (24). Briefly, the genes had their TAT signal peptides removed from the sequences as determined by SignalP6.0 (35). The synthetic genes were inserted into the *p15TV-L* plasmid linearized with BseRI digestion using Gibson assembly.

Expression plasmids for point mutants CfrA-Y80W, CfrA-F125W, CfrA-C260Y, CfrA-Y256F, and double mutants CfrA-Y80W-C260Y and CfrA-Y80W-Y256F were constructed using the site-directed PCR mutagenesis protocol from Edelheit *et al.* (36). The wild-type CfrA plasmid, *p15TVL-cfrA*, was used as the template for mutagenesis for the single mutants, and the constructed *p15TVL-Y80W* plasmid was used as the template for the double mutants. Primers used to introduce the mutations are in Table S1. All assembled plasmids were confirmed by Sanger sequencing using universal T7 and T7 terminator primers. CfrA-Y80W-F125W was codon optimized and synthesized in the *pET-21* plasmid along with sequences encoding a ribosome binding site, N-terminal histidine tag, and TEV cleavage site added upstream of the gene (Twist Bioscience). Additional mutants and enzymes are described in Table S2 and their activity can be found in the supplementary material.

All expression plasmids were co-transformed with *pBAD42-BtuCEDFB* into *E. coli* BL21(DE3) *cat-araC-*P_BAD_-*suf*, Δ*iscR::kan*, Δ*himA::*Tet^R^, abbreviated to *E. coli ara-Suf* Δ*iscR* (37, 38). The *pBAD42-BtuCEDFB* plasmid, used for cobalamin uptake, was generously provided by the Booker Lab (Pennsylvania State University, PA, USA). *E. coli* BL21(DE3) *cat-araC*-P_BAD_-*suf*, *ΔiscR::kan, ΔhimA::*Tet^R^, used for enhanced iron-sulfur production, was generously provided by the Antony and Kiley Labs (St. Louis University School of Medicine, MO, USA; University of Wisconsin-Madison, WI, USA).

### Expression and Nickel-Affinity Chromatography

Methods for RDase expression and enrichment by nickel-affinity chromatography have been described elsewhere (20, 24). Minor amendments were made here. Briefly, for each enzyme, an overnight starter culture was used to inoculate 1 L, 500 mL, or 250 mL of Luria Broth (LB) in 2 L, 1 L, or 500 mL media bottles, respectively, to ensure consistent liquid and headspace volumes. The cultures were grown aerobically at 37°C, 170 rpm until the O.D._600_ reached 0.2–0.4, at which 3 µM hydroxycobalamin and 0.2% (w/v) ʟ-arabinose were supplemented to induce Btu expression. The cultures were capped and incubated again at 37°C until the O.D._600_ reached ∼0.6–0.8, after which the cultures were purged with N_2_ gas for 1 hr/L liquid volume. The cultures were supplemented with 50 µM cysteine and 50 µM ammonium ferric citrate, and RDase expression was induced with 0.3–0.5 mM isopropyl β-ᴅ-1-thiogalactopyranoside (IPTG). Finally, the culture was incubated for 18 hr at 15°C, 170 rpm.

The cultures were kept anaerobic and pelleted by centrifugation, and all subsequent steps were performed in the glovebox or sealed vessel to maintain anaerobicity. The pellet was resuspended in 5 mL of Lysis Buffer (50 mM Tris-HCl pH 7.5, 150 mM NaCl, 0.1% Triton, 5% glycerol) per 1 g of wet cell weight; a 1 L culture would usually yield 3-4 g. Reagents 10x Bug Buster concentrate (used at 1x), 1 mM tris(2-carboxyethyl)phosphine (TCEP), 50 µg/mL leupeptin, 2 µg/mL aprotinin, 0.3 mg/mL lysozyme, 1.5 µg/mL DNase, and 10 mM MgCl_2_ (final concentrations) were all added fresh. This was incubated at room temperature shaking at 50 rpm for 20 min. The lysate was clarified by centrifugation at 30 000 xg, 4°C for 20 min and incubated with nickel-nitrilotriacetic acid (Ni-NTA) beads either in the column for lysate quantities ≥ 10 mL, or in batch for smaller lysate quantities.

The Ni-NTA beads were washed with Wash buffer (50 mM Tris-HCl pH 7.5, 150 mM NaCl, 30 mM imidazole) until no protein could be detected with Bradford reagent. The RDases were eluted using Elution buffer (50 mM Tris-HCl pH 7.5, 150 mM NaCl, 300 mM imidazole). The eluant was exchanged into Storage buffer (50 mM Tris-HCl pH 7.5, 150 mM NaCl) and concentrated using a 30 kDa cut-off Millipore filter tube. The protein was stored in liquid nitrogen. The concentrate was quantified with a Bradford assay and the purity was assessed by SDS-PAGE using Image Lab v6.1 Software (Bio-Rad Laboratories Inc., 2020; Figure S1 and Table S5 in the accompanying Excel).

### Reductive Dechlorination Assays

The dechlorination activity of each mutant and wild-type enzyme was assessed on the substrates chloroform (CF), 1,1,1-trichloroethane (TCA), 1,1,2-TCA, and 1,1-dichloroethane (DCA); some enzymes were also assessed for activity on 1,2-DCA and dichloromethane (DCM) (Figure S4). Each enzyme assay was carried out anaerobically in 500 µL reaction volumes using 2-mL vials with 500 µL glass inserts sealed with a Teflon coated cap such that there was no headspace in the vial. The reaction mix contained 5 mM Ti(III) citrate and 2 mM methyl viologen in 50 mM Tris-HCl pH 7.5. Substrates were added to a target concentration of 0.5 mM using saturated water stocks. Enzymes were added in a volume of 5 µL to deliver 0.2–0.6 µg of RDase (accounting for purity). The reactions were incubated at room temperature for 65 min. The reaction was stopped by transferring 400 µL into 5.6 mL HCl-acidified water (pH < 2) in an 11-mL headspace vial and crimped for analysis by gas chromatography-flame ionization detection (GC-FID).

The samples were all analyzed using by GC-FID separation on an Agilent 7890A GC instrument equipped with an Agilent GS-Q plot column (30 m length, 0.53 mm diameter). Sample vials were equilibrated to 70°C for 40 min in an Agilent G1888 autosampler, then 3 mL of the sample headspace was injected (injector set to 200°C) onto the column by a packed inlet. The flow rate was 11 mL/min of helium as the carrier gas. The oven was held at 35°C for 1.5 min, raised at a rate of 15°C/min up to 100°C, the ramp rate was reduced to 5°C/min until the oven reached 185°C at which point it was held for 10 min. Finally, the oven was ramped up to 200°C at a rate of 20°C/min and held for 10 min. The detector was set to a temperature of 250°C. Data were analyzed using Agilent ChemStation Rev. B.04.02 SP1 software, and peak areas were converted to liquid concentration using external standard curves for each compound.

Enzyme-free and free cobalamin (0.0005 mg) negative control reactions were performed for each substrate. All reactions were done in technical triplicates; CfrA and DcrA were assayed using four replicates. The mass of the dechlorinated product was calculated from the liquid concentration and apparent activity was calculated by dividing the product mass by the incubation time and the amount of enzyme in the reaction. All data for OG 97 RDases and CfrA mutants are in Figures S2 and S3, additional activity values on DCM and 1,2-DCA are shown in Figure S4. All raw data for activity is reported in the accompanying Excel document in Tables S6 and S7.

### Activity Normalization

The substrate preference of each enzyme was visualized using a bubble plot. For each enzyme, the activity level on substrates CF, 1,1,1-TCA, and 1,1-DCA was normalized by dividing by the highest activity observed by that particular enzyme (Equation 1 in Supplemental Text S3). This normalization scaled the activities from 0 (no activity) to 1 (highest activity). The size of each bubble in the plot represents the normalized activity, allowing for a visual comparison of how each enzyme interacts with the three substrates. However, it does not allow for a direct comparison of activity levels between different enzymes. Additionally, in Text S3 a dendrogram was produced to cluster the enzymes based on activity similarity (Figure S5). This clustering was performed by normalizing the activity of all enzymes to the activity of CfrA and DcrA depending on the substrate (see Equations 2 and 3 and Text S3 for further details).

### Protein Models and Docking

New protein models of CfrA, DcrA, and TmrA were obtained through the AlphaFill webserver with cobalamin and [4Fe-4S] clusters incorporated into the structures (39). The models were retrieved using UniProt IDs: CfrA, K4LFB7; DcrA, J7I1Z7; TmrA, T0I1B4. The models were relaxed using the YASARA energy reduction webserver (40). Water molecules were removed from these models and they were used for docking studies with AutoDock4.2 (41). Each protein was docked with CF, 1,1,1-TCA, 1,1-DCA, and 1,1,2-TCA to proximate the binding location and residue interactions with each substrate. All AutoDock settings were kept as default, cobalt was added to the parameters file as a heteroatom, the enzymes were kept rigid with no flexible residues, and the docking grid space was located on the open face of the docked cobalamin molecule. The top docking position for each enzyme-ligand pairing was used to assess binding affinity and interacting side chains (Table S3).

Protein models of AcdA and the mutant RDases were produced using AlphaFold2 (33). The models were superimposed with the cobalamin and [4Fe-4S] clusters from the CfrA model in PyMOL and exported as a new model to incorporate the co-factors. The edited models were relaxed using the YASARA energy minimization server. These were all docked with CF, 1,1,1-TCA, 1,1-DCA, and 1,1,2-TCA using the same methods as described above. Images of the undocked protein structure active sites are in Figures S6 and S7.

The steric clash between Tyr256 rotamers and the surrounding residues was assessed using the Wizard mutagenesis tool in PyMOL. The relative distance between mutated residues was also recorded to compare which mutations introduce more strain in the model. The observed strain score and the rotamer probability are in Table S4 and Figures S8 and S9.

## Results and Discussion

### Active Site Transplant

Previous studies probing the roles of specific residues have focused on PceA_S_ and enzymes that target haloethenes (26–28, 31), yet many of the key residues established in these studies are variable within OG 97 and specifically between CfrA and DcrA. When structurally modelled, it is apparent that the differences between CfrA and DcrA are localized in the active site, particularly in the substrate access channel and binding pocket (Figure 1A and B). Phillips *et al.* suggested five residues within the putative active site that may influence the activity and the mechanism of these chloroalkane reductases (34). Three of these residues at positions 80, 260, and 391, align structurally with Phe38, Tyr246, and Trp376 of PceA_S_, respectively. Residues 262 and 388 were also proposed to have functional significance in OG 97, but residue 262 is identical in both CfrA and DcrA so it will not be explored here (34). Additionally, the mutation of Trp118 in DcaA from *D. dichloroeliminans* has been shown to have a detrimental effect on its activity, and the equivalent residues 125 in CfrA and DcrA diverge (28). Therefore, in this study we have chosen five variable active site residues to mutate in CfrA and DcrA based on previous analyses and proposed roles: residues 80, 125, 260, 388, and 391. All of these residues differ between CfrA and DcrA, but the rest of OG 97 members have varying combinations of these residues, which might contribute to their more flexible substrate selection (Table 3). A full alignment of OG 97 enzymes is in Supplemental Text S1.

**Figure 1.**
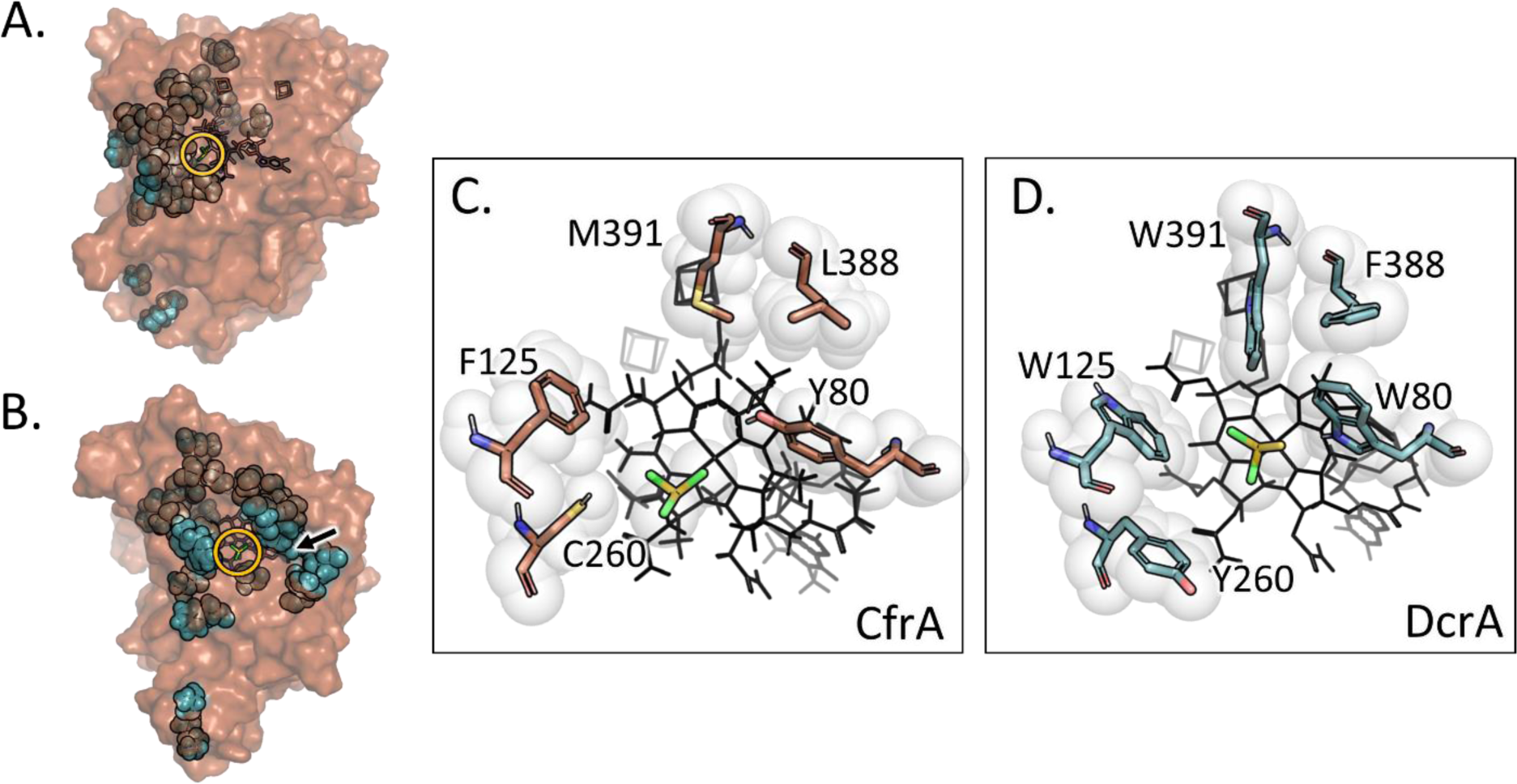
Surface model of CfrA in orange with the amino acids that differ from DcrA shown as blue spheres and chloroform docked in the active site. (A) Profile and (B) face view of the active site. The yellow circle indicates the location of the substrate. The black arrow indicates the substrate access channel. The five targeted active site residues in (C) CfrA (orange) docked with chloroform and (D) DcrA (blue) docked with 1,1-dichloroethane. Substrates are shown with yellow backbones and cobalamin and [4Fe-4S] cofactors are shown as black wire structures.

**Table 2.** Enzyme and plasmid description for RDase mutants discussed in the main text.

| Enzyme Expressed | Plasmid | Description |
| --- | --- | --- |
| CfrA | *p15TVL-cfrA* | Wild-type CfrA |
| DcrA | *p15TVL-dcrA* | Wild-type DcrA |
| CfrA-5M | *p15TVL-cfrA-5M* | CfrA with mutations: Y80W, F125W, C260Y, L388F, and M391W |
| DcrA-5M | *p15TVL-dcrA-5M* | DcrA with mutations: W80Y, W125F, Y260C, F288L, and W391M |
| CfrA-3M | *p15TVL-cfrA-3M* | CfrA with mutations: C260Y, L388F, and M391W |
| CfrA-Y80W-F125W | *pET21-cfrA-Y80W-F125W* | CfrA with mutations: Y80W and F125W |
| CfrA-Y80W-Y256F | *p15TVL-cfrA-Y80W-Y256F* | CfrA with mutations: Y80W and Y256F |
| CfrA-Y80W-C260Y | *p15TVL-cfrA-Y80W-C260Y* | CfrA with mutations: Y80W, C260Y |
| CfrA-F125W | *p15TVL-cfrA-F125W* | CfrA with F125W mutation |
| CfrA-Y80W | *p15TVL-cfrA-Y80W* | CfrA with Y80W mutation |
| CfrA-Y256F | *p15TVL-cfrA-Y256F* | CfrA with Y256F mutation |
| CfrA-C260Y | *p15TVL-cfrA-C260Y* | CfrA with C260Y mutation |

**Table 3.**
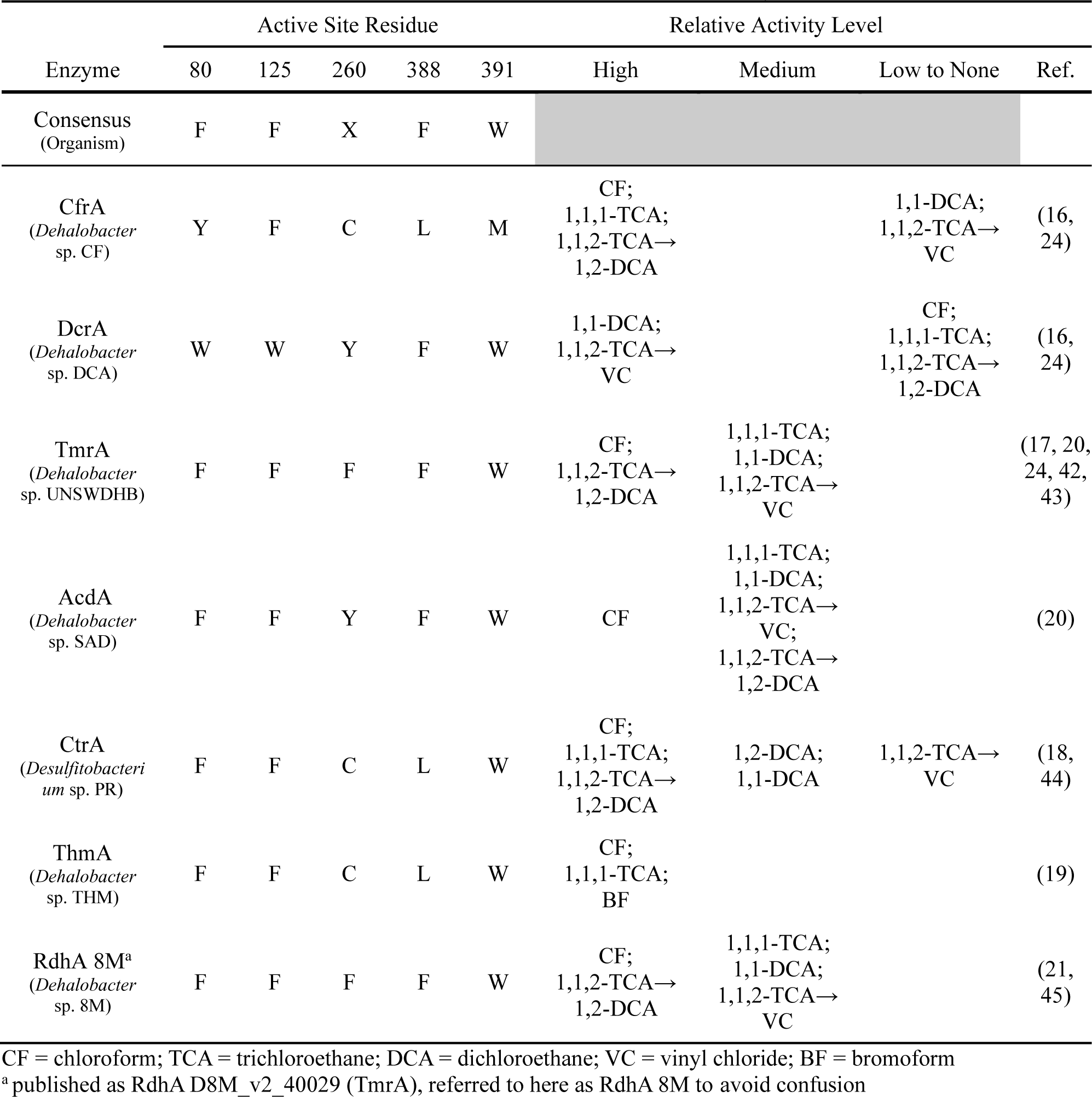
Active site residues in all OG 97 members and their relative activity on known substrates.

We conducted site-directed mutagenesis with all five residues of interest switched to those of the opposite enzyme, i.e., CfrA was mutated to have the same five residues of DcrA and vice versa. Effectively, mutants CfrA-5M and DcrA-5M have been transplanted with the active site of the other while all other structural differences remained untouched. These mutants were used to determine if the substrate binding site is the sole determinant of the wild-type enzymes’ specificity. The roles of specific residues and their interactions were further analyzed in CfrA with subsets of the original five mutations. These included CfrA-3M—a variant targeting the three aliphatic residues in CfrA—and several double mutants and point mutants (all described in Table 2). The wild-type enzyme and mutants were all expressed in *E. coli* and extracted using nickel-affinity chromatography. The purity of the enzymes ranged due to variations in solubility and expression levels, so the protein concentrations were adjusted to account for purity (all protein values are in Table S5 in the accompanying Excel and Supplemental Text S2). In general, DcrA and its mutants had low solubility and purity; thus, the mutagenesis studies were primarily carried out using CfrA variants. Information on the DcrA mutants can be found in Supplemental Text S5.

### Mutants Shift Substrate Specificity

As expected, the CfrA-5M mutant did indeed display DcrA-like behaviour with diminished activity on CF and 1,1,1-TCA compared to wild-type CfrA, and a 10-fold increase in activity on 1,1-DCA (Figure 2). However, CfrA-5M still has a 1.1 nmol/s/mg greater turnover rate of CF than 1,1-DCA. Further, the level of activity on 1,1-DCA only reached a little over a third of the activity displayed by DcrA. These findings affirm that while the five amino acids targeted within the active site do influence the substrate selectivity, but additional differentiating factors may also contribute to enhanced activity. While this study focuses on active site disparities, potential additional sources of differentiation will be explored later. Here we continue to dissect the role that these five residues play in the gain of activity on 1,1-DCA.

**Figure 2.**
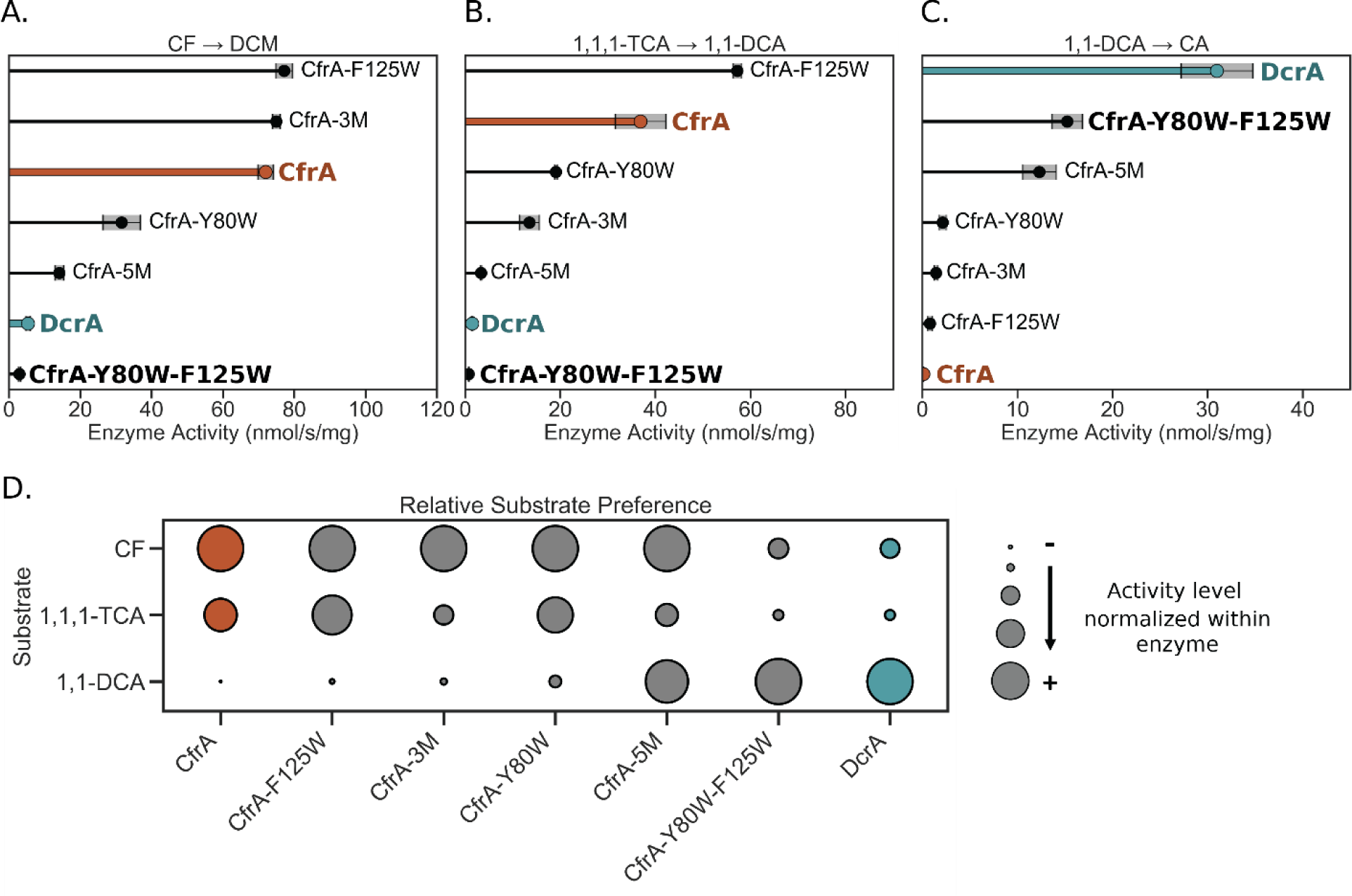
Dechlorination activity of CfrA, DcrA, and CfrA mutants on (A) chloroform, (B) 1,1,1-trichloroethane, and (C) 1,1-dichloroethane. CfrA activity is highlighted in orange and DcrA in blue. Grey shading indicates the standard deviation between samples (n = 3, n = 4 for CfrA and DcrA). (D) The activity of each enzyme normalized to the substrate they have the highest activity on to show their relative preferences, larger circles indicate higher relative activity levels. CF = chloroform, DCM = dichloromethane, TCA = trichloroethane, DCA = dichloroethane, CA = chloroethane.

After confirming that transplanting the active site of DcrA into CfrA caused over a 100-fold increase in the dechlorination of 1,1-DCA by CfrA-5M, we sought to determine whether this change is due to the role of an individual residue or a compounding effect by several residues. Structural analysis of the five targeted residues in the wild-type RDases shows that DcrA has a much more compact substrate binding pocket than CfrA (Figure 1C and D). DcrA has many large aromatic residues that add steric bulk and may help to stabilize 1,1-DCA binding but consequently prevent larger substrates like CF from binding efficiently. CfrA, on the other hand, has three small aliphatic residues (cysteine, leucine, and methionine) that create space to accommodate the trichlorinated substrates. The compact active sites of PceA_D_ and DcaA were suggested to be essential for the correct positioning of the substrates—mutations that widened the active sites were disruptive to chloroethene dehalogenation (28). To narrow down the contributions of different amino acids, we first targeted CfrA’s aliphatic residues Cys260, Leu388, and Met391—which are all aromatic in DcrA—in the subsequent mutant CfrA-3M.

Despite CfrA-3M containing the most divergent mutations between CfrA, DcrA, and the rest of OG 97, the activity of CfrA-3M was not very different from the wild-type CfrA (Figure 2). The activity of CfrA-3M on CF was unchanged but its activity on 1,1,1-TCA decreased by half, suggesting some steric interference with the additional methyl group on the substrate. There was a 10-fold increase in activity on 1,1-DCA; however, this is marginal compared to CfrA-5M. Given these results, we chose to target the two remaining residues, Tyr80 and Phe125, to assess their role in activity on 1,1-DCA.

Figure 2 Both single and double mutants targeting Tyr80 and Phe125 were assayed for activity against CF, 1,1,1-TCA, and 1,1-DCA. The double mutant CfrA-Y80W-F125W exhibited the most profound change in activity shifting substrate preference to be more similar to that of DcrA. Figure 2D depicts the substrate profile of each enzyme variant, where activity is normalized to the substrate with the highest activity for that enzyme. The substitution of only two residues having a profound effect on substrate preferences was not expected. These two changes in CfrA-Y80W-F125W caused an almost complete loss in activity on CF and 1,1,1-TCA and display how finely tuned substrate selection can be in RDases. Although CfrA-Y80W-F125W gained a 130-fold level of activity on 1,1-DCA, its activity was still only half that of DcrA (Figure 2C). 1,1-DCA has a lower reduction potential (E_0_’ = 397 mV) than CF (560 mV) and 1,1,1-TCA (561 mV) (46), which means that there is less of a driving force for the reduction of 1,1-DCA if the same electron donor is used. To make this reaction more energetically favourable, DcrA may have peripheral modifications that alter the redox potential of the cobalamin or [4Fe-4S] cofactors. In particular, [4Fe-4S]-dependent enzymes have been shown to use the amino acids that surround the clusters to tune the cluster redox potential for optimal activity (47–49). We explored some potential mutants targeting residues near the cofactors in DcrA, but these yielded limited changes in activity (discussed in Supplemental Text S5).

Interestingly, CfrA-F125W produced a 1.5-fold increase in turnover of 1,1,1-TCA and a similar turnover of CF as CfrA (Figure 2). Conversely, CfrA-Y80W resulted in diminished activity on both CF and 1,1,1-TCA, both resulting in approximately half the wild-type activity. It is clear that the introduction of Trp80 has a more adverse effect on CfrA’s activity on trihalogenated substrates. But neither mutant showed increases in 1,1-DCA reduction to the level of CfrA-Y80W-F125W, indicating that these residues must work cooperatively to produce this activity. This further supports the idea that smaller substrates require a tighter active site to be held in the ideal conformation.

### Mutants Change Mechanism Preference

In addition to the changes in substrate selectivity observed by CfrA-5M and CfrA-Y80W-F125W, we also observed a shift in the transformation pathway of 1,1,2-TCA. Members of OG 97 can carry out both the hydrogenolysis of 1,1,2-TCA to 1,2-DCA or the dihaloelimination to vinyl chloride (VC) (Figure 3C), but the ratio of products varies depending on the enzyme (17, 24, 44, 45). CfrA and DcrA are both on the extreme ends of either mechanism, with each producing almost homogenous products of either 1,2-DCA by CfrA or VC by DcrA (Figure 3).

**Figure 3.**
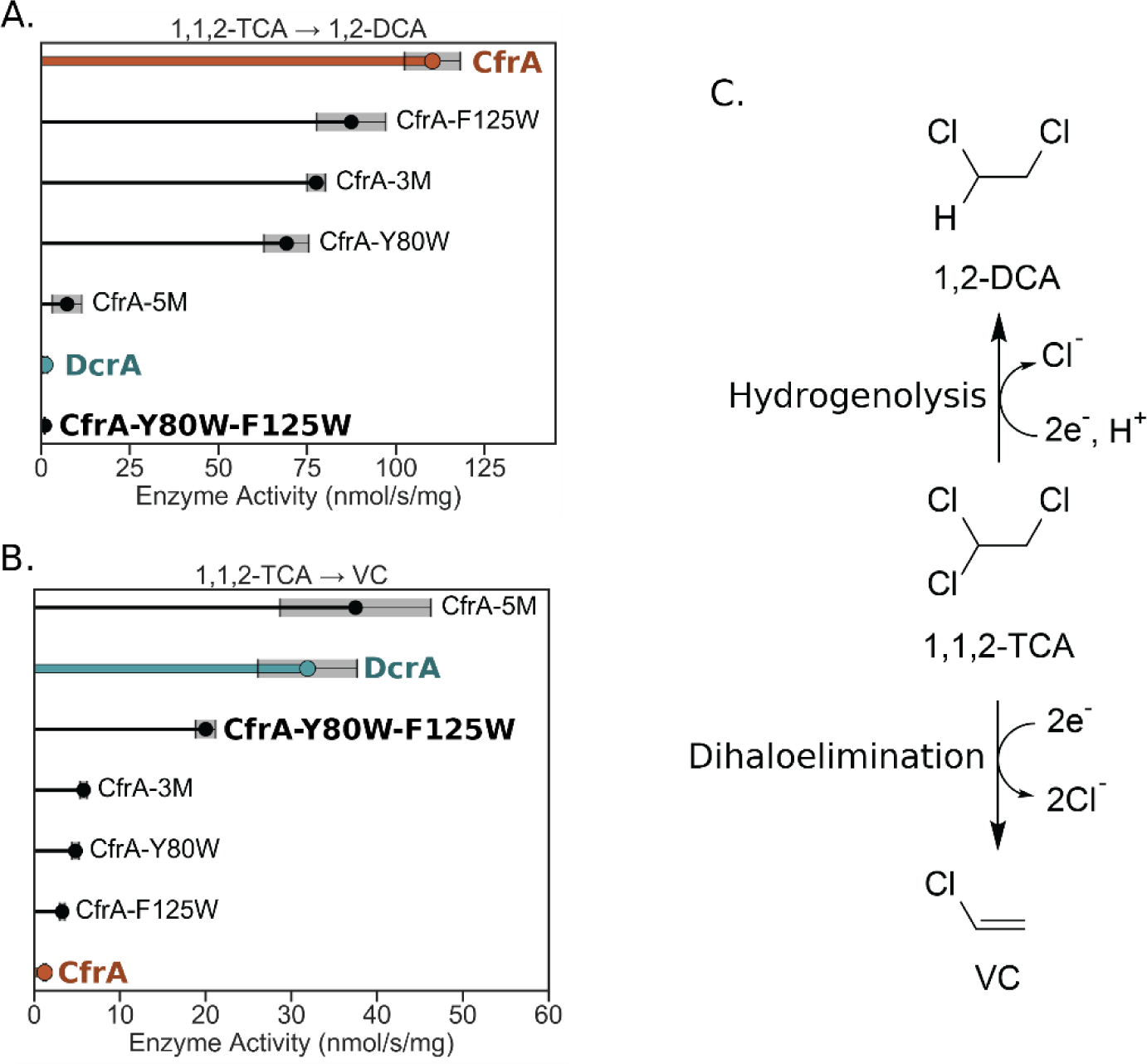
Dechlorination activity of CfrA, DcrA, and mutants on 1,1,2-trichloroethane to produce (A) 1,2-dichloroethane or (B) vinyl chloride. (C) The two reaction pathways that 1,1,2-trichloroethane can undergo. CfrA activity is highlighted in orange and DcrA in blue. Grey shading indicates the standard deviation between samples (n = 3, n = 4 for CfrA and DcrA). TCA = trichloroethane, DCA = dichloroethane, VC = vinyl chloride.

CfrA-5M and CfrA-Y80W-F125W both shifted to produce a majority of VC from 1,1,2-TCA, akin to DcrA. Once again, the CfrA-3M, CfrA-Y80W, and CfrA-F125W mutants show similar reaction profiles to CfrA, with a high preference for hydrogenolysis (>90% 1,2-DCA product, Figure 3). The dihaloelimination reaction of 1,1,2-TCA is thermodynamically favoured with a reduction potential 201 mV higher than the hydrogenolysis reaction (46, 50). Given the greater driving force for reducing 1,1,2-TCA to VC, CfrA must create interactions with the substrate that enable the hydrogenolysis pathway. However, given that the point mutants do not have an appreciable shift towards the elimination reaction, CfrA most likely uses a network of interactions to stabilize its substrates. When these interactions are disrupted, the enzyme will revert to the easier elimination reaction as we saw in CfrA-5M and CfrA-Y80W-F125W.

When considering multiple dichlorination pathways with regards to remediation, the transformation of 1,1,2-TCA to 1,2-DCA is the preferred pathway not only because VC is ranked higher and more toxic on the Substance Priority List by the Agency for Toxic Substances and Disease Registry (1), but also because VC strongly inhibits CfrA dechlorination activity on CF and 1,1,1-TCA (51). Understanding how structural variants affect the substrates and products of RDases is crucial for predicting outcomes in bioremediation processes. However, as we have demonstrated here, this is not a trivial task.

### Substrate-Enzyme Interaction Observations

Interactions between residues and their substrates, as well as inter-residue interactions, are highly complex and often challenging to predict, especially when relying on static structural models. By employing protein docking and visualizing the active site, we can make informed predictions about the interactions that could be disrupted by mutagenesis, thereby leading to changes in activity. However, it is important to consider the limitations of current structure prediction tools, particularly regarding the precise rotamers and locations of side chains within the structure. Thus, we will be discussing our general observations of potential structural differences based on the chemical characteristics of the mutated residues.

The two major differences between DcrA and CfrA are the additional steric bulk in DcrA due to the presence of more aromatic residues and the positioning of residues that can act as hydrogen bond donors. The mutations made in CfrA-5M were all from smaller amino acids to larger aromatic residues. These residues not only take up space but can also provide hydrophobic interactions with the substrates. Yet, we observed that only two of those mutations were necessary to achieve activity similar to DcrA. Residues Tyr80 and Phe125 line the substrate access channel and mouth of the active site in the CfrA model (Figure 4A). Mutating both of these residues to tryptophan could narrow the channel. Both CF and 1,1,1-TCA bear three chlorine atoms on one carbon. Since chlorine has a much larger atomic radius than hydrogen, the widest point of these substrates will be larger than 1,1-DCA or 1,1,2-TCA. The steric restriction of the channel may occlude the binding of these larger substrates. In the mutagenesis of *D. dichloroeliminans* DcaA, the opposite effect was observed, where decreasing the bulk at the equivalent of Phe125 (Trp432 in DcaA) decreased the activity on all substrates (28). The data reported here, however, suggest that residue 80 has a greater impact on enzyme activity. CfrA-Y80W consistently has lower activity than CfrA, but CfrA-F125W actually has increased activity on 1,1,1-TCA.

**Figure 4.**
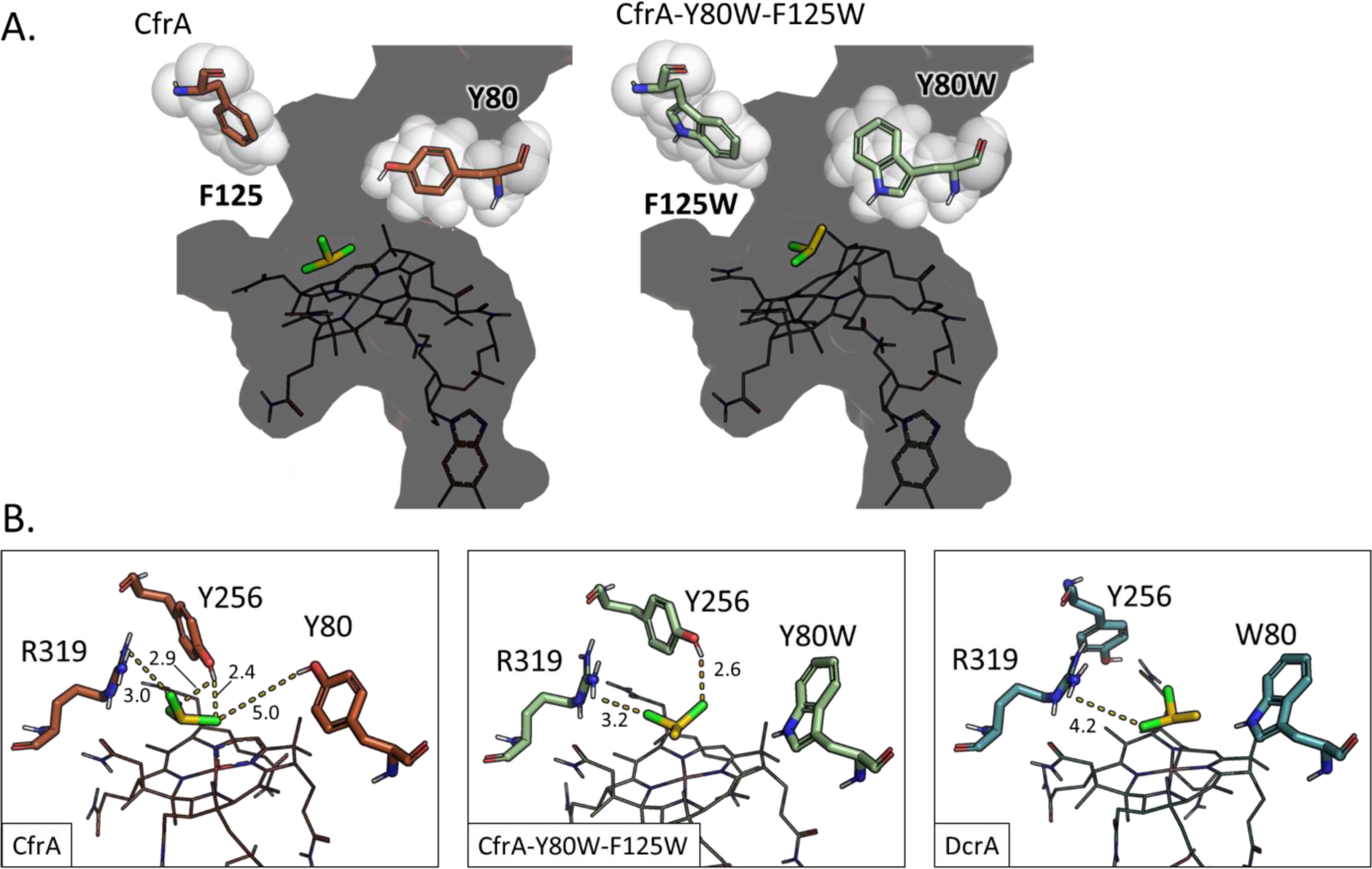
Protein structure models of CfrA, CfrA-Y80W-F125W, and DcrA show the possible interactions made by residues 80 and 125. (A) The positioning of residues 80 and 125 in the substrate access channel of CfrA (left; orange) and CfrA-Y80W-F125W (right; green). (B) Potential hydrogen-bond donors in the active sites of CfrA (orange), CfrA-Y80W-F125W (green), and DcrA (blue). Distances of atoms are labelled in Å on dashed lines. All models are docked with their preferred substrate (yellow): CfrA with chloroform, CfrA-Y80W-F125W and DcrA with 1,1-dichloroethane.

Another differentiator between CfrA and CfrA-Y80W-F125W is the potential for hydrogen (or halogen) bonding by Tyr80 and other neighbouring residues. Tyrosine forms stronger hydrogen bonds and has a higher propensity to form halogen bonds than tryptophan (52–54). The hydroxyl group has the ability to create a strong bond with the chlorines located on the substrate, which may help in directing the substrate into a reactive orientation. While Tyr80 appears far from the substrate in the docked model, there is a high degree of uncertainty as to the exact catalytic docking site and position of the residue. Additionally, the interaction could occur upon entry through the channel. The DcrA model showed putative interactions between Trp80 and docked substrates and could provide more hydrophobic stabilization (Table S3). Therefore, the number and type of stabilizing interactions CfrA and DcrA make with their substrates could be a factor in their selectivity.

In the CfrA active site, there are three potential hydrogen bond donors—Tyr80, Tyr256, and Arg319 (Figure 4B). Arg319 is predicted to be the proton donor involved in catalysis and is structurally conserved amongst all OG 97 members (31). Tyr256 is conserved in all OG 97 sequences, but its predicted positioning in the protein structures varies (Figure S9). When the models were docked with their substrates, CfrA was the only one which had Tyr256 interacting with the substrates (Table S3). While we cannot be certain of this residue’s orientation in the models, we can look at how the surrounding residues may interact with it.

In Figure 5 the relative positions of Tyr256 are shown in relation to the varying residues 125 and 260 in CfrA and DcrA. DcrA has bulky aromatics at both of these positions, and thus, steric clash may prevent Tyr256 from rotating into the active site, while CfrA does not have these restraints. To illustrate the potential steric clash and assess how our designed mutations may affect the Tyr256 positioning, we analyzed the different Tyr256 rotamers in the CfrA-5M model using the PyMOL Wizard mutagenesis tool. Using this tool, we are able to assess possible rotamers of Tyr256 and how likely they are to have steric clash with neighbouring residues. In Figure 5, the steric clash is shown with red disks, for which the size and density of the disks show the relative degree of overlap between the van der Waals radii of the atoms in the residues—i.e. greater overlap and steric clash displays more red disks in the images and would correspond to a less favourable conformation. In the CfrA-5M model active site, there is much less room for Tyr256 and a lot of steric clash is predicted if it faces toward the substrate binding location (Figure 5C and D). Both F125W and C260Y contribute to this clash, as we still see steric overlap when the mutations are visualized individually (Figure 5E and F), but the preferred rotamers for the single mutants are not as obvious. Thus, the steric clash between the introduced mutations and other important active site residues could also contribute to the activity changes we see in the CfrA mutants (Table S4).

**Figure 5.**
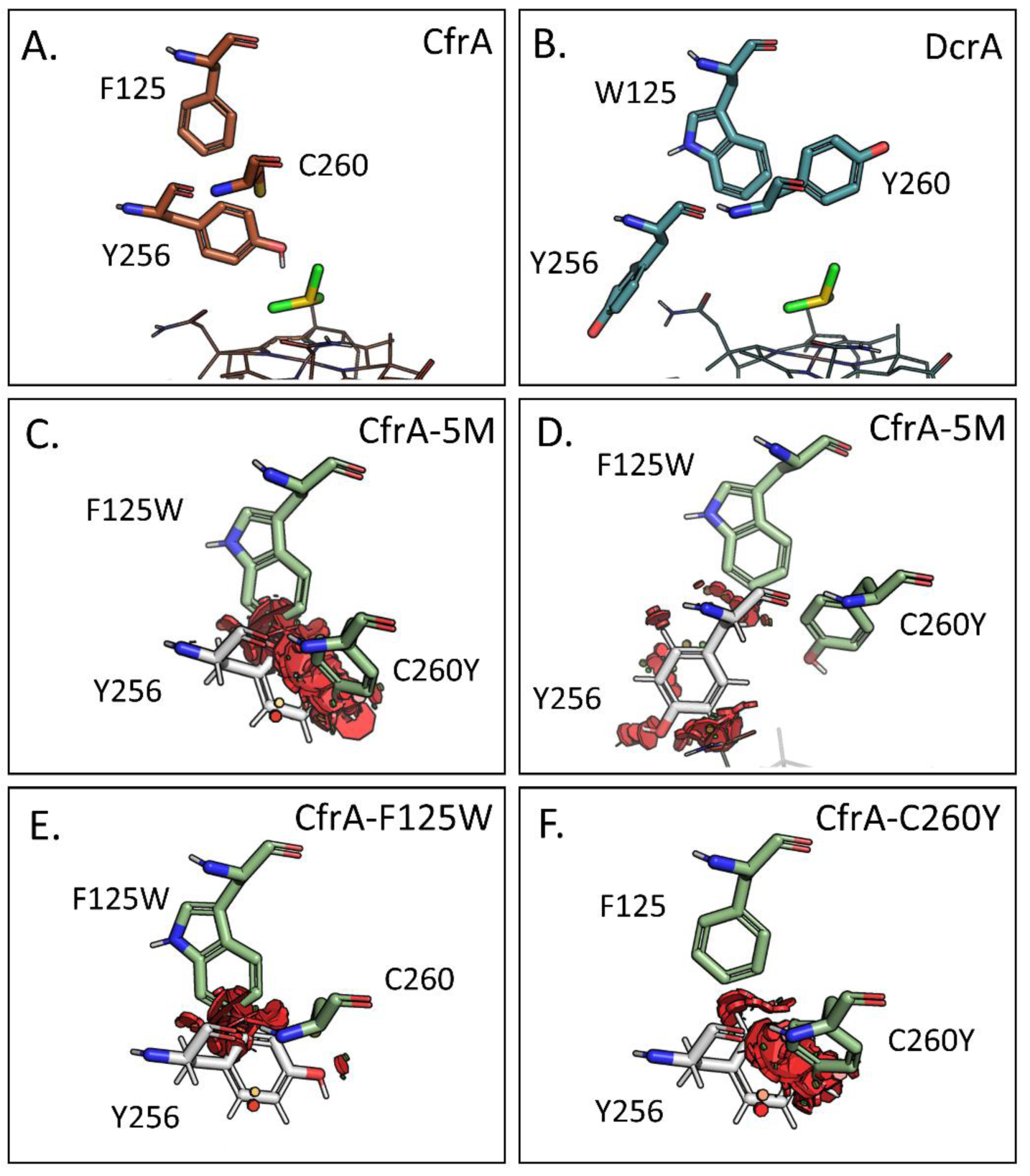
Protein structure models of (A) CfrA and (B) DcrA show the predicted positioning of Tyr256 in the active site; the docked substrate is shown in yellow and cobalamin is shown as a wire figure. CfrA-5M model with the steric clash resulting from the (C) Tyr256 (white) rotamer pointing toward the active site and (D) the rotamer pointing out of the active site. Tyr256 pointing toward the active site and the resulting clash with each of the (E) F125W and (F) C260Y mutations individually. Red disks indicate steric clash where Tyr256 overlap with the van der Waals radius of atoms in neighbouring residues; only residues at positions 125 and 260 are shown as green sticks.

It is unclear whether Tyr256 is important in substrate selectivity in OG 97, but the equivalent Thr294 in DcaA was shown to have a great impact on substrate selectivity (28). The T294V mutant of DcaA enhanced preference for dehalogenation of TCE over PCE and reduced dihaloelimination activity on 1,1,2-TCA and 1,2-DCA (28). The authors attributed this change to a slight increase in residue size, thereby suggesting that dihaloelimination may be favoured when there is more space at this position. This could be mirroring what we observe between CfrA, DcrA, and the mutants. Another possibility is the potential stabilizing hydrogen bonding between Tyr256 and the substrate. To delve into the possible interactions of Tyr256 more, we produced several more double and single mutations targeting the steric clash and hydrogen bonding potential.

### Influence of Individual Residues

The second round of mutants was produced to assess the role of Tyr256 and Cys260, particularly in 1,1,2-TCA transformation. To determine if the hydrogen bonding capability of Tyr256 is important to activity, it was mutated to phenylalanine. This change would still maintain the bulk and hydrophobicity provided by the aromatic ring, with only a small decrease in size. Cys260 was targeted due to its proximity to Tyr256, as it may clash with Tyr256 and push it out of the active site (Figure 5F). Further, residue 260 is highly variable within OG 97, and Tyr260 is present in DcrA and AcdA, which both perform the dihaloelimination reaction more readily than CfrA (20, 24). Both the Y256F and C260Y were paired with Y80W in double mutants of CfrA, because Y80W appeared to have more influence over activity than F125W alone.

The single mutants (Y256F and C260Y) did not hold much control over the substrate preferences of CfrA (Figure 6). It is unlikely that Tyr256 participates in hydrogen bonding for substrate selection because it maintained the same activity level and preferences as CfrA (Figure 6 and Figure 7). Yet, when paired with Y80W, the CfrA-Y80W-Y256F double mutant lost 60-70% of its activity on CF, 1,1,1-TCA and 1,1,2-TCA. A similar trend was seen with CfrA-C260Y, where it retained activity on CF and 1,1,1-TCA but lost 70% of the activity when paired with Y80W.

**Figure 6.**
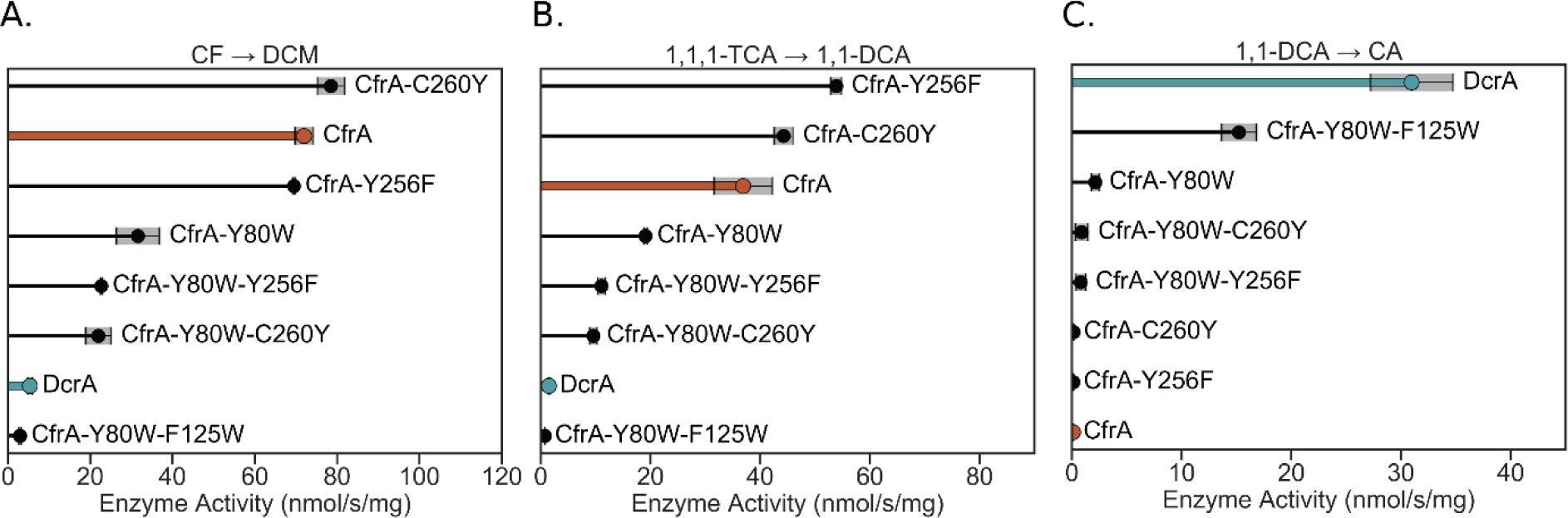
Dechlorination activity of CfrA, DcrA, and select mutants on (A) chloroform, (B) 1,1,1-trichloroethane, and (C) 1,1-dichloroethane. CfrA activity is highlighted in orange and DcrA in blue. Grey shading indicates the standard deviation between samples (n = 3, n = 4 for CfrA and DcrA). CF = chloroform, DCM = dichloromethane, TCA = trichloroethane, DCA = dichloroethane, CA = chloroethane.

**Figure 7.**
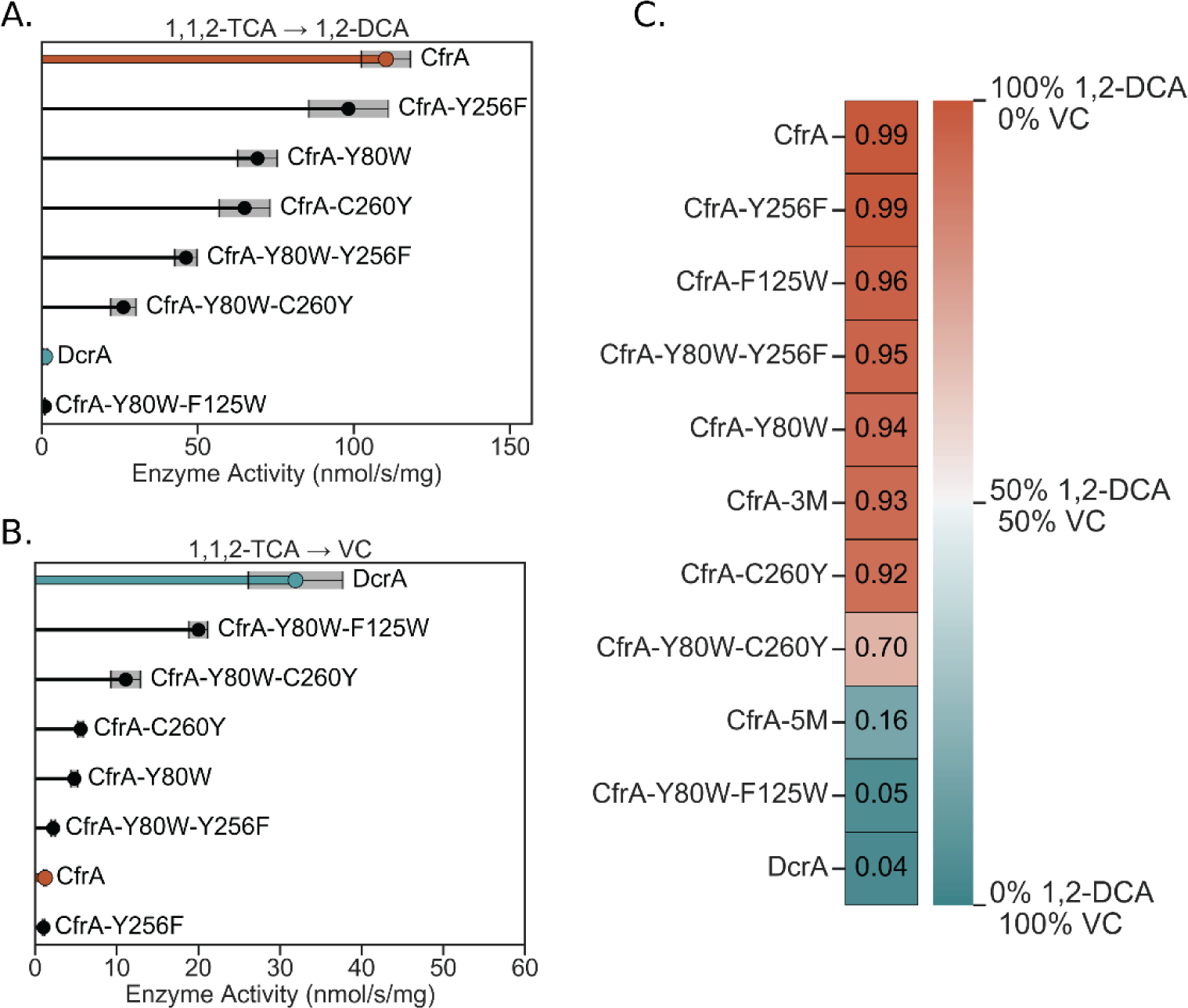
Dechlorination activity of CfrA, DcrA, and select mutants on 1,1,2-trichloroethane to produce (A) 1,2-dichloroethane or (B) vinyl chloride. (C) The proportion of the dechlorination product that went to 1,2-dichloroethane (more orange) and vinyl chloride (more blue); the number in the square indicates the ratio of 1,2-dichloroethane product. CfrA activity is highlighted in red, DcrA in blue, and CfrA-Y80W-F125W in green. Grey shading indicates the standard deviation between samples (n = 3, n = 4 for CfrA and DcrA). TCA = trichloroethane, DCA = dichloroethane, VC = vinyl chloride.

Neither CfrA-Y80W-Y256F nor CfrA-Y80W-C260Y could reduce 1,1-DCA. This supports the idea that increasing the bulk at the substrate access channel, as we see in CfrA-Y80W-F125W, improves activity on smaller and less halogenated substrates. Perhaps narrowing the substrate access channel forces the substrate into a more catalytically active position and prevents it from leaving before catalysis. Kinetic analysis to assess how residues 80 and 125 impact the *K_M_* on 1,1-DCA would be needed to further support this hypothesis.

While the change in Tyr256 did not affect the transformation pathway of 1,1,2-TCA, the C260Y mutation did have a slight effect, especially when paired with Y80W. CfrA-Y80W-C260Y retained less than 25% of the wild-type hydrogenolysis activity on 1,1,2-TCA but increased the dihaloelimination reaction 10-fold (Figure 7). On its own, the C260Y mutation shifted the percentage of VC in the product from 1% (wild-type) to 8%, and the double mutant CfrA-Y80W-C260Y increased the percent VC in the product to 30% (Figure 7C). The large increase in VC production from the double mutant further indicates that there are multiple interactions by CfrA to suppress dihaloelimination. The effect of C260Y may be due to its bulk preventing Tyr256 from entering the active site, and the open space increases the dihaloelimination activity, as we have discussed. Tyr80, on the other hand, could be involved in changing the orientation of 1,1,2-TCA as it enters the active site and could also provide stabilization through hydrogen bonding. TmrA and AcdA have both a non-hydrogen-bonding Phe80 and bulky aromatic residues at 260 which could be why they produce mixtures of VC and 1,2-DCA (Table 3 and Figure S3). Experimental structures of CfrA and DcrA are needed to confirm the positioning of Tyr256 and support its role in activity divergence.

### Significance and Conclusions

By manipulating the amino acid composition of CfrA’s active site we have found residues that are highly influential in selecting larger trichlorinated substrates versus dichlorinated, and those that can direct the reaction mechanism of 1,1,2-TCA toward one product over another. The two mutations that completely changed CfrA’s substrate specificity were Y80W and F125W. These residues line the entrance to the active site from the substrate access channel and may act as a filter to block larger trihalogenated substrates while also stabilizing 1,1-DCA for catalysis. Corresponding residues have been reported or suggested to be important in determining the activity of PceA_S_, PceA_D_, and DcaA due to their proximity to the active site (3, 26–28). Similar to DcaA, we observed that added bulk in the active site has an influence on the size of the preferred substrates.

Further, we have observed that the active site residues in OG 97 produce compounding effects on enzyme activity and substrate selectivity. No single point mutation increased 1,1-DCA activity; however, residue 80 seems to have the most influence in selecting trihalogenated substrates. When several active site mutations were combined, we saw the greatest increase in non-wild-type activity, suggesting synergistic contributions to function.

By analyzing the diverging evolution of CfrA and DcrA that specialized them toward separate substrates, and comparing this to chloroethene-reducing RDases, we have identified several residues that could be hotspots for natural evolution and protein engineering. Changes in residues 80, 125, and 256 (Phe38, Trp96, Thr242 in PceA_S_), both in the side chain and potentially the positioning of the residues, appear to have large impacts on the preferred substrate size and the tendency to perform dihaloelimination (28). The primary factor that is consistent between the results from CfrA and DcaA mutagenesis is the steric bulk of the residues. The addition of bulkier residues, particularly at the mouth of the active site, favours smaller substrates. Introducing more space into the active site has been detrimental to the activity on small substrates, but there is no clear trend on the active site space and inclination for dihaloelimination (28). Variations between residues might also alter a network of conserved residues, so surrounding residues should also be considered when introducing mutations. Crystal structures and molecular dynamics analysis of CfrA and DcrA would help confirm these observations.

It is unlikely that the active site is the only source of substrate differentiation, and other areas that should be further assessed are residues at the opening and along the substrate access channel. As shown in Figure 1 the natural variations between CfrA and DcrA are clustered around the substrate channel; key differences here could be why the CfrA mutants do not reach the same level of activity on 1,1,-DCA as the wild-type DcrA. Another impact on enzyme activity—which has yet to be explored in RDases—is the environment surrounding the iron-sulfur clusters and cobalamin. It is known that the protein structure impacts the reduction potential of redox-active cofactors; factors such as solvation, and hydrogen-bonding or ionic residues surrounding the iron-sulfur clusters can drastically change their reduction potential and therefore, the enzyme’s activity (49, 55).

Additionally, the findings suggest that CfrA interacts differently with its substrates, CF and 1,1,1-TCA, using distinct active site residues. The mutants, particularly CfrA-3M, showed varied changes in activity levels with each substrate. Adding steric bulk in the active site reduced 1,1,1-TCA activity more than CF, while changes in the substrate access channel impacted both similarly. Kinetic analysis indicates that CfrA has a higher *K_M_* with 1,1,1-TCA (51, 56, 57), meaning lower affinity. This difference in *K_M_* and sensitivity to active site size aligns with CfrA isotope fractionation data showing increased masking (less fractionation) of 1,1,1-TCA compared to CF (58–60). Masking occurs when the rate is governed by a non-fractionating slow step—i.e. substrate binding—rather than the transformation step. Members of OG 97 have varying levels of masking when using CF; CfrA has the least masking and has fractionation similar to cobalamin (34, 45, 60). The considerable effect that minor active site differences can have on substrate selection shown here, suggests that the masking likely results from differences in substrate binding. However, further isotope analyses and mechanistic studies are needed to confirm.

The findings presented here underscore the challenge of accurately predicting RDase activity based solely on sequence information. One cannot presume identical activity between RDases, even if they share high sequence similarity. Additionally, they contribute to our understanding of the factors influencing substrate selectivity in RDases and hold promise for guiding future rational engineering efforts to accommodate novel substrates.

## Acknowledgments

This work was funded by the Natural Science and Engineering Research Council (NSERC) through a Discovery Grant and Canada Research Chair to E.A.E., a Canadian Graduate Scholarship – Doctoral to K.J.P. and an Undergraduate Student Research Award to C.B. Funding was also provided by the Ontario Government with an Ontario Graduate Scholarship to K.J.P. The funders had no role in the study design, data collection and interpretation, or the decision to submit the work for publication.

We would like to thank the Booker Lab from Pennsylvania State University for gifting *the pBAD42-BtuCEDFB* plasmid and the Antony Lab from the St. Louis University School of Medicine and the Kiley Lab from the University of Wisconsin-Madison for gifting us the *E. coli* BL21(DE3) *cat-araC*-P_BAD_-*suf* Δ*iscR*::*kan* Δ*himA*::Tet^R^ strain. We also thank Peter Stogios and Sofia Lemak for their input on the structural analyses and manuscript.

E.A.E. and K.P. conceived the project. C.B. contributed to the cloning, purification, and preliminary activity assays of the first round of mutants. All other experiments and data analysis were performed by K.P. The manuscript was written by K.P. and E.A.E. All co-authors contributed to manuscript revision and have approved the submitted version.

